# An efficient system for production of highly purified inhibins using mammalian cell expression and tandem affinity purification

**DOI:** 10.64898/2026.09.10.750725

**Authors:** Evan R. S. Buddle, Cynthia S. Hinck, Yeu-Farn Lin, Ying Wang, Troy C. Krzysiak, Daniel J. Bernard, Andrew P. Hinck

**Affiliations:** Department of Pharmacology and Therapeutics, McGill University, Montreal, Québec, Canada; Department of Structural Biology, University of Pittsburgh School of Medicine, Pittsburgh, PA 15260, USA; Department of Anatomy and Cell Biology, McGill University, Montreal, Québec, Canada

**Keywords:** inhibin, protein purification, reproduction, structure-function, endocrinology, pituitary gland

## Abstract

Inhibin A and B (InhA and InhB) are potent negative regulators of follicle-stimulating hormone (FSH), which is essential for female fertility. InhA and InhB are heterodimeric transforming growth factor β (TGF-β) family ligands consisting of an inhibin α (inhα) subunit disulfide linked to an inhibin βA or βB (inhβA, inhβB) subunit, respectively. During purification, inhibins must be separated from activins (inhβA or inhβB homodimers), which always form when producing inhibins, and have the opposite effect on FSH synthesis. Current inhibin purification methods are technically challenging, require reagents that are not generally available, result in low purity or contamination with activins, or employ substitutions within the growth factor domain to favor inhibin formation over activin. We developed a simple and high-yielding expression and purification system for inhibins by transfection of suspension-cultured mammalian cells and tandem affinity chromatography, followed by reverse-phase purification of the mature ligands. We purified 1.4 mg (n = 3, SD = 0.47) and 0.093 mg (n = 3, SD = 0.043) of wild-type InhA and InhB per liter of culture medium with little to no contaminating activin A or B, at equal to higher purity and bioactivity than commercially available inhibins. Thus, we present a simple, robust, and versatile system for producing inhibins to interrogate their functions and mechanisms of action *in vitro* and *in vivo*.

## Introduction

Inhibin A and B (InhA and InhB) are transforming growth factor β (TGF-β) family ligands that are produced in the gonads and act at the level of the pituitary to inhibit follicle-stimulating hormone (FSH) synthesis (1, 2). FSH is critical for female fertility and its production from pituitary gonadotrope cells is positively regulated by hypothalamic gonadotropin-releasing hormone and activin-class TGF-β family ligands, such as myostatin, growth differentiation factor 11 (GDF11), and activin A (3–7). FSH consists of an α subunit shared with other dimeric pituitary glycoproteins, and a unique β subunit encoded by *Fshb/FSHB* (1). Activin-class ligands stimulate, whereas inhibins decrease *Fshb/FSHB* expression.

TGF-β family ligands are produced as larger precursor proteins that are proteolytically cleaved by furin family proprotein convertases, resulting in a pro-domain that remains non-covalently associated with the mature growth factor (GF) domain. The pro-domain may or may not confer latency (2, 8). GFs are comprised of two disulfide-linked ‘cystine knot’ protomers, each of which resemble an ‘open hand’, with an α-helix forming the wrist and β-strands forming fingers that interact with type I and type II serine/threonine kinase receptors to exert biological activity (2).

Pro-InhA and Pro-InhB are synthesized as heterodimers composed of the inhibin α (inhα) subunit connected by a single disulfide bond, within the GF domain, to either an inhibin βA or βB (inhβA, inhβB) subunit, respectively (2, 9). The pro-protein heterodimers are cleaved at the consensus furin cleavage site between the pro- and GF-domains in both the α- and β-subunits, thus separating the GF heterodimers from their component pro-domains. The α-subunit is also cleaved at a secondary furin site near the N-terminus of the pro-domain (2, 10). Pro-ActA and pro-ActB are synthesized as inhβA or inhβB homodimers (2, 10–13). Activin-class ligands induce FSH production by binding two type II activin receptors, ACVR2A or ACVR2B, and then bind and activate two type I activin or TGF-β receptors (5, 14–16). Activated type I receptors phosphorylate SMAD2/3, which associate with SMAD4 and accumulate in the nucleus to act with FOXL2 to promote *Fshb/FSHB* transcription, the rate-limiting step in FSH production (7, 17–20).

Inhibins do not bind type I receptors and thus lack intrinsic signaling activity (2, 21). They function instead as antagonists of activin-class ligands by sequestering the type II receptors, ACVR2A and ACVR2B (5, 21). Inhibins bind non-signaling type III receptors, primarily through their α-subunit. Together with type II receptor binding through their single β-subunit, this enables sufficient avidity for inhibins to compete for type II receptors against activin-class homodimers (2, 22–24). The zona pellucida C (ZPC) domain of TGF-β receptor III (TGFBR3 or betaglycan) binds InhA and InhB, with greater affinity for the former (22, 23). TGFBR3-like (TGFBR3L), an ortholog of the TGFBR3 ZPC domain expressed exclusively in pituitary gonadotropes, is specific for InhB (23, 25, 26). It is unknown how InhA or InhB bind their co-receptors or the structural basis that explains co-receptor selectivity. Betaglycan ZPC also binds to TGF-βs. Insights from structural studies of TGF-β:betaglycan complexes indicate that the fingers of the inhα subunit may be involved in co-receptor binding (27, 28). To address these questions, inhibins must be produced in sufficient yield and purity for structure-function analyses, which has been historically challenging.

The predominant challenge for production of recombinant inhibins is complete removal of co-produced activins, which have opposite actions (29, 30). It is also necessary to remove any pro-domains that may be present (2, 10). Procedures for producing InhA and InhB using stably transfected mammalian cells or co-transfection of plasmids encoding the inhibin α and β subunits in mammalian cells have been described (29, 31, 32). Resulting proteins were assessed via western blotting and functional assays (29, 31, 32). The inhibins purified from stably transfected cells are reliable and of high purity (31). However, their purification requires up to four orthogonal purification steps, including, in some cases, affinity chromatography with the proprietary anti-inhα (“R1”) antibody (31, 32). Lastly, the cells are not generally available and the inhibins produced are wild-type alone.

The transfection-based method described by Goney and co-workers allows for production of both wild-type protein and variants, thus providing greater flexibility (29). However, the system as published, while including tags for purification, was mostly directed toward generating inhibins for cell-culture assays without or with only partial purification of the ligands from the conditioned medium. To enable this, the subunits were modified to: 1) disfavor inhibin β subunit homodimerization (inhβA^A347H^, inhβB^G329E^), 2) enhance inhibin potency (inhα^S334I^), 3) decrease potency of contaminating activins (inhβA^A347H^, inhβA^M334A^, inhβB^G329E^, inhβB^M402A^), and 4) improve processing at the secondary furin cleavage site (super-cut, or ‘SCUT’) in the α-subunit (29). This is not suitable for structural and biophysical studies, which require fully purified wild-type InhA and InhB. This method of inhibin production may also be flawed for use in reporter assays, as it assumes reduced activin activity and limited production.

## Results

### Design of bis-cistronic plasmids to produce wild-type and mutant InhA

Initially, we followed the procedure described by Goney and co-workers (29) and designed InhA constructs, designated as version 1 (V1), that contained the specified amino acid modifications (**Fig. 1A-B**) (29). We also used a dual-tagging scheme similar to that described by Goney (29), with a nona-histidine (his) tag on inhα inserted just upstream of the primary furin processing site, and a Strep-tag II affinity (hereafter, strep) tag, rather than the Flag tag as in (29), on the N-terminus of the inhβA pro-domain. We also used orthogonal factor Xa processing sites instead of the endogenous furin processing sites at the pro-GF boundary, as furin processing is often incomplete and can complicate purification. We also introduced a G316A substitution in the βA-subunit to eliminate a possible secondary factor Xa processing site near the N-terminus of the mature βA-subunit.

**Figure 1:**
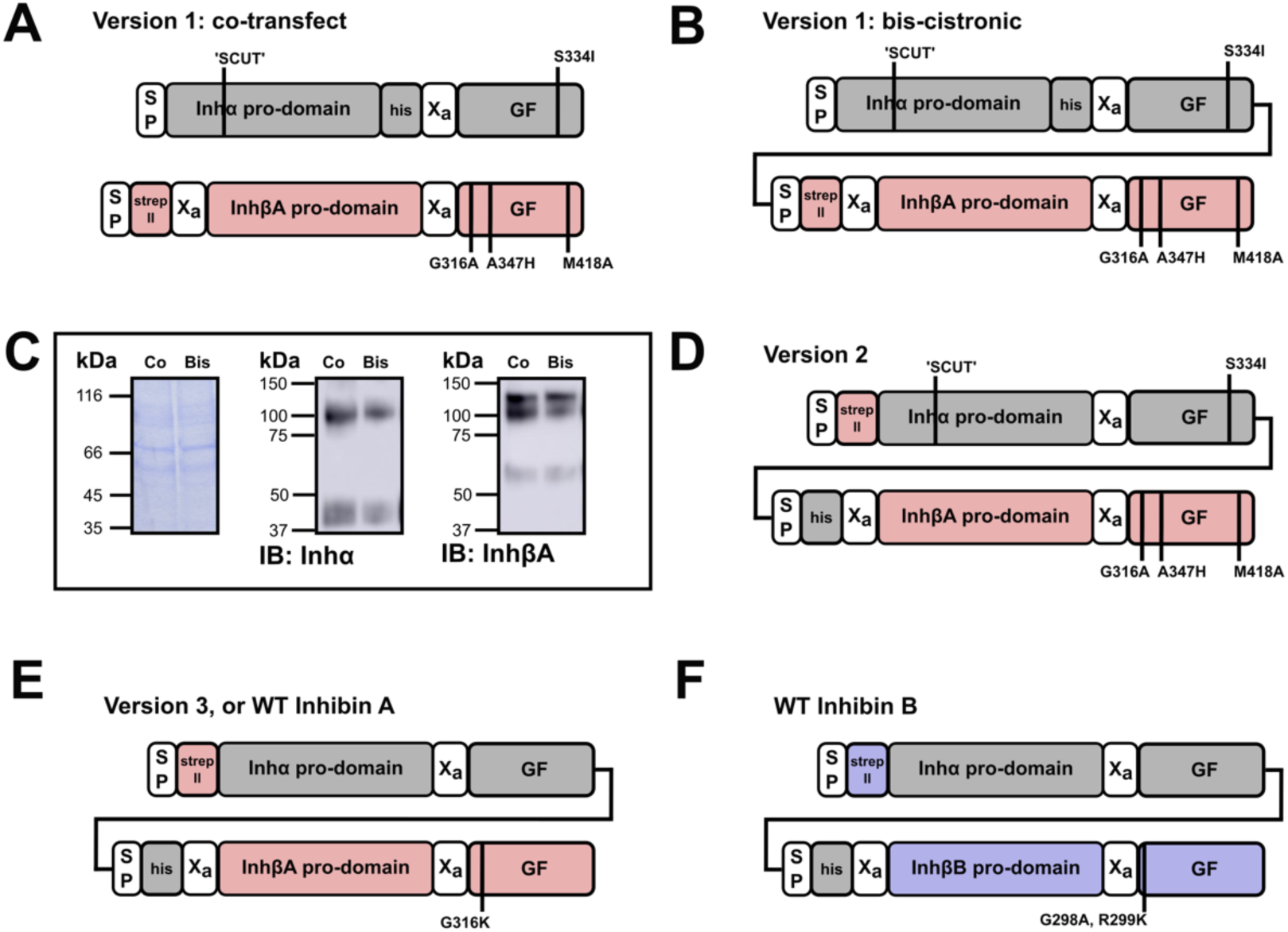
Constructs for inhibin production in mammalian cells. **(A)** Plasmid design for production of ActA (V1) and InhA (V1) by co-transfection, based on the design proposed by Goney et al., 2020. Relevant modifications to the coding sequences are marked and labelled. The signal peptide (SP) and factor Xa (Xa) sites are boxed in white. **(B)** Bis-cistronic plasmid design for producing InhA V1. **(C)** Coomassie-stained SDS-PAGE gel (27.5 μL/lane; left panel) and immunoblots (5 μL/lane; middle and right panels) of conditioned media (CM) from Expi293 cells co-transfected with individual inhα and inhβA subunits (panel **A**) or transfected with a single bis-cistronic InhA V1 construct (panel **B**). (**D)** Bis-cistronic plasmid design for InhA (V2), in which the positions of the strep and his tags were swapped relative to V1. **(E)** Bis-cistronic plasmid design for wild-type InhA (V3 or WT). This plasmid includes the modification inhβA^G316K^ to eliminate a secondary factor Xa site. **(F)** Bis-cistronic plasmid design for wild-type inhibin B, with inhβB^G298A^ and inhβB^R299K^ to eliminate a secondary factor Xa site.

We first tested production of pro-InhA V1 protein from cells co-transfected with inhα and inhβA plasmids (**Fig. 1A**) compared to transfection of a single inh*a*:inhβA bis-cistronic plasmid (**Fig. 1B**). The overall amount of protein, as well as the ratio of α to βA, produced by the two methods was similar (**Fig. 1C**). In some cases, as shown in **Fig. 1C**, the overall amounts obtained by co-transfection were slightly higher (33), but this was not consistent. Therefore, to simplify the procedure, we proceeded with transfection of a single bis-cistronic plasmid. Subsequently, optimized bis-cistronic plasmids were designed to produce other versions of InhA or InhB (**Fig. 1D-F**).

### Pro-ActA binds non-specifically to cobalt immobilized metal affinity chromatography columns

To assess whether we could scale up production of InhA V1, we transfected Expi293 suspension-adapted HEK293 cells with an empty vector (EV) construct, the ActA (mono-cistronic inhβA) V1, or the InhA V1 bis-cistronic construct described above (**Fig. 1A-B**, **Fig. 2A**). Upon loading and eluting the EV-conditioned medium (CM) onto the cobalt immobilized metal affinity chromatography (Co-IMAC) column, non-specific proteins eluted between 14–26 %B along the 0 to 0.5 M imidazole gradient (**Fig. 2A**). There was no inhα or inhβA immunoreactivity detected in the flow through and wash or eluted fractions (**Fig. 2B-C, Supp. Fig. S1A**). Upon loading the pro-ActA-CM onto the Co-IMAC column and eluting with the same imidazole gradient, nearly all of the pro-ActA bound to the column (**Fig. 2B, lane 2, Supp. Fig. S1B)** and was present in the eluted fractions, predominantly between 14-34 %B. This was unexpected as the inhβA V1 construct lacks a his-tag (**Fig. 2D-G, lanes 2-7)**. Only after immunoblotting against a large excess of CM did we detect pro-ActA in the combined flow-through and wash from the Co-IMAC (**Fig. 2B, lane 8**). Upon loading the pro-InhA CM, it behaved similarly, though eluted at higher imidazole concentrations, 21-49 %B, which was expected given the inclusion of a his-tag on the α-subunit (**Fig. 2E, 2H-I, lanes 3-10)**. Despite eluting at a higher imidazole concentration compared to pro-ActA, there was still significant overlap of pro-ActA in the pro-InhA eluate (**Fig. 2H-I, lanes 4-10**). In addition, these blots demonstrated that the inhβA^A347H^ mutation did not fully impair pro-ActA dimerization (**Fig. 2G, lanes 2-10**), although the observed inhβA monomer may reflect an impairment in its ability to form dimers (**Fig. 2D-E, 2G, 2I, lanes 2-10**).

**Figure 2:**
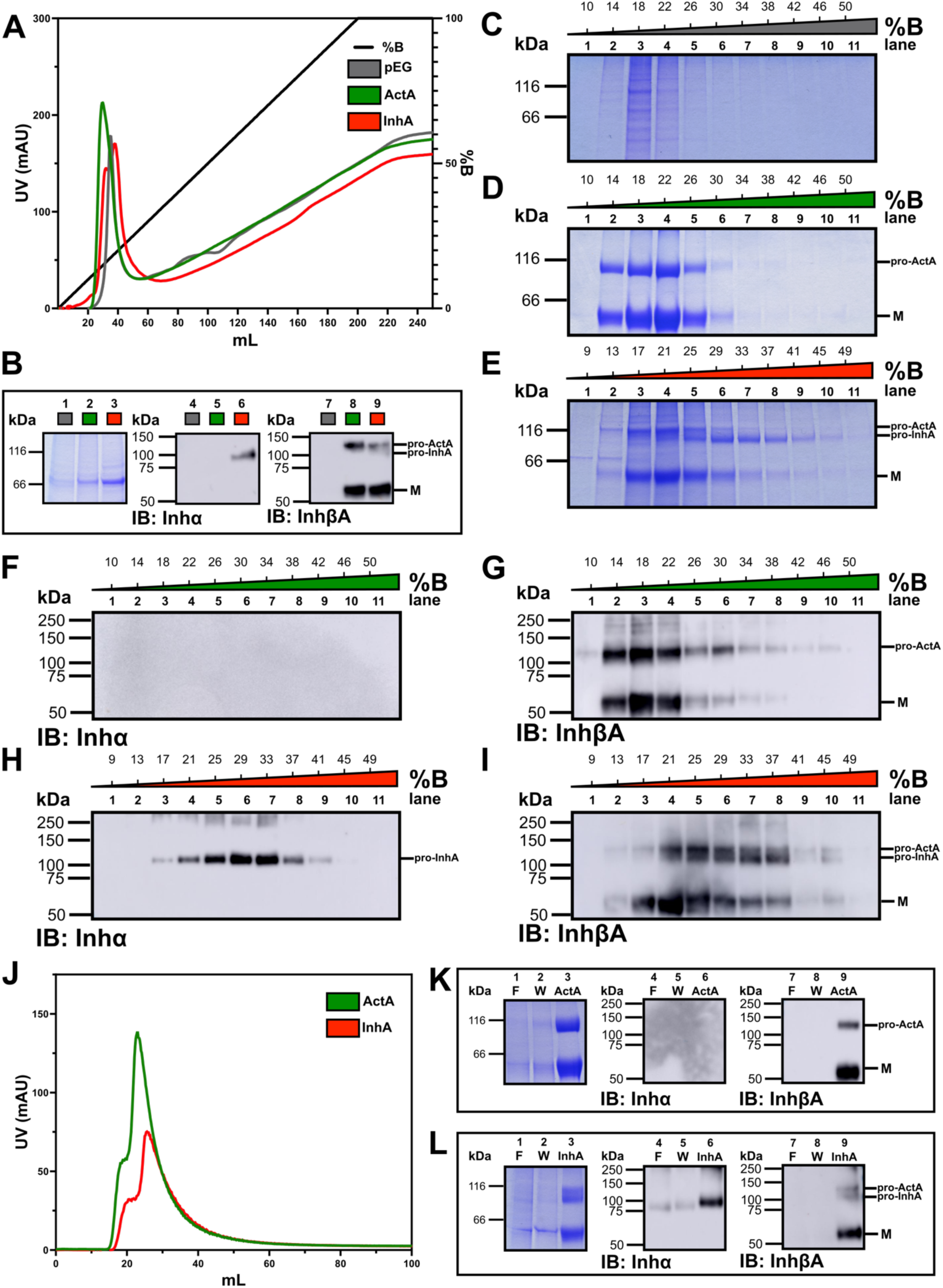
ActA binds non-specifically to the cobalt immobilized metal affinity chromatography (Co-IMAC) column. (**A)** CM from Expi293 cells transfected with empty vector (pEG) or constructs to produce pro-ActA V1 or pro-InhA V1 was loaded onto a Co-IMAC column. Protein was eluted and fractionated over a 200 mL, 0-0.5 M imidazole gradient (%B). Traces were aligned to the start of the gradient. **(B)** Non-reducing SDS-PAGE analysis with Coomassie-staining (left) and immunoblots (middle and right) corresponding to combined flow-through and wash from the Co-IMAC purification for pEG, pro-ActA, and pro-InhA (flow-through and wash were concentrated to 80 mL prior to analysis). For the SDS-PAGE gels in **(B)**, 27.5 μL/lane of sample was used for Coomassie staining, and 10 μL/lane was used for immunoblotting. Coomassie-stained SDS-PAGE gel (non-reducing) of collected fractions (27.5 μL/lane) corresponding to eluted protein for **(C)** pEG, **(D)** pro-ActA, and **(E)** pro-InhA, with markers denoting pro-ActA or pro-InhA dimers or monomer (M). Immunoblots (non-reducing) against the inhα or inhβA subunit for collected fractions (0.55 μL/lane) from the Co-IMAC purification of **(F-G)** pro-ActA and **(H-I)** pro-InhA. **(J)** Chromatograms of biotin-eluted pro-ActA and pro-InhA from the strep column (traces were aligned to the start of elution). Non-reducing SDS-PAGE gels stained with Coomassie (left) and immunoblots (middle and right) corresponding to flow-through (F), wash (W), and eluted protein (concentrated to 1 mL) from the strep affinity purification of **(K)** pro-ActA and **(L)** pro-InhA. For SDS-PAGE corresponding to strep purification steps, in this figure and Fig. 4F, 5F, 6F, 2 μL/lane of sample was used for Coomassie-staining and 1 μL/lane of a 1/25 dilution was used for immunoblotting.

Protein-containing fractions from the Co-IMAC purified pro-ActA and pro-InhA were pooled and purified using a Strep-Tactin XT affinity column (hereafter strep column) (**Fig. 2J**). As expected, the strep-tagged pro-ActA bound to the column, as nothing was detected in the flow-through or wash (**Fig. 2K, lanes 1-2, 7-8**), and pro-ActA was eluted from the column (**Fig. 2K, lanes 3, 9**). In the purification of pro-InhA V1, anti-inhα immunoblotting revealed a small amount of InhA did not bind (**Fig. 2L, lanes 4-5**), but because the strep tag resided on the inhβA subunit, most of the protein was in the eluate, as shown by strong staining for both pro-ActA and pro-InhA (**Fig. 2L, lanes 3, 6, 9)**.

### Separation of ActA and InhA with the V1 InhA construct using reverse phase high-performance liquid chromatography

Following the tandem purification of the expressed pro-ActA and pro-InhA, we cleaved the purified proteins with factor Xa and used reversed-phase high-performance liquid chromatography (RP-HPLC) to separate the GFs from the pro-domains, or fragments thereof. Using an optimized elution gradient from 10 to 50% acetonitrile (%B) over 30 column volumes, ActA GF (peak ‘b’) eluted at 33-33.4 %B (**Fig. 3A-J**), while InhA GF (peak ‘d’) eluted at 35-37 %B (**Fig. 3B, E-F, I-J**). InhA (peak ‘d’) contained both glycoforms (31- and 34-kDa) of the mature GF, consistent with earlier reports (2) (**Fig. 3E-F, I-J, peak ‘d’, lanes 5-6**). RP-HPLC purification effectively separated the InhA and ActA GFs, as no band corresponding to ActA (**Fig. 3J, peak ‘b’, lane 3)** was detected in the InhA fractions (**Fig. 3I-J, peak ‘d’, lanes 5-6**) by immunoblotting against the inhβA subunit. However, there were some higher molecular weight (HMW) species observable by SDS-PAGE analysis and detected by immunoblotting, which may correspond to fragments of the pro-domain that were not fully cleaved, thus co-eluting with the InhA GF (**Fig. 3E-F, 3I-J, peak ‘d’, lanes 5-6**). The amounts of InhA and ActA obtained from expression of the InhA V1 construct in this purification were approximately equal, at around 0.25 mg per liter.

**Figure 3:**
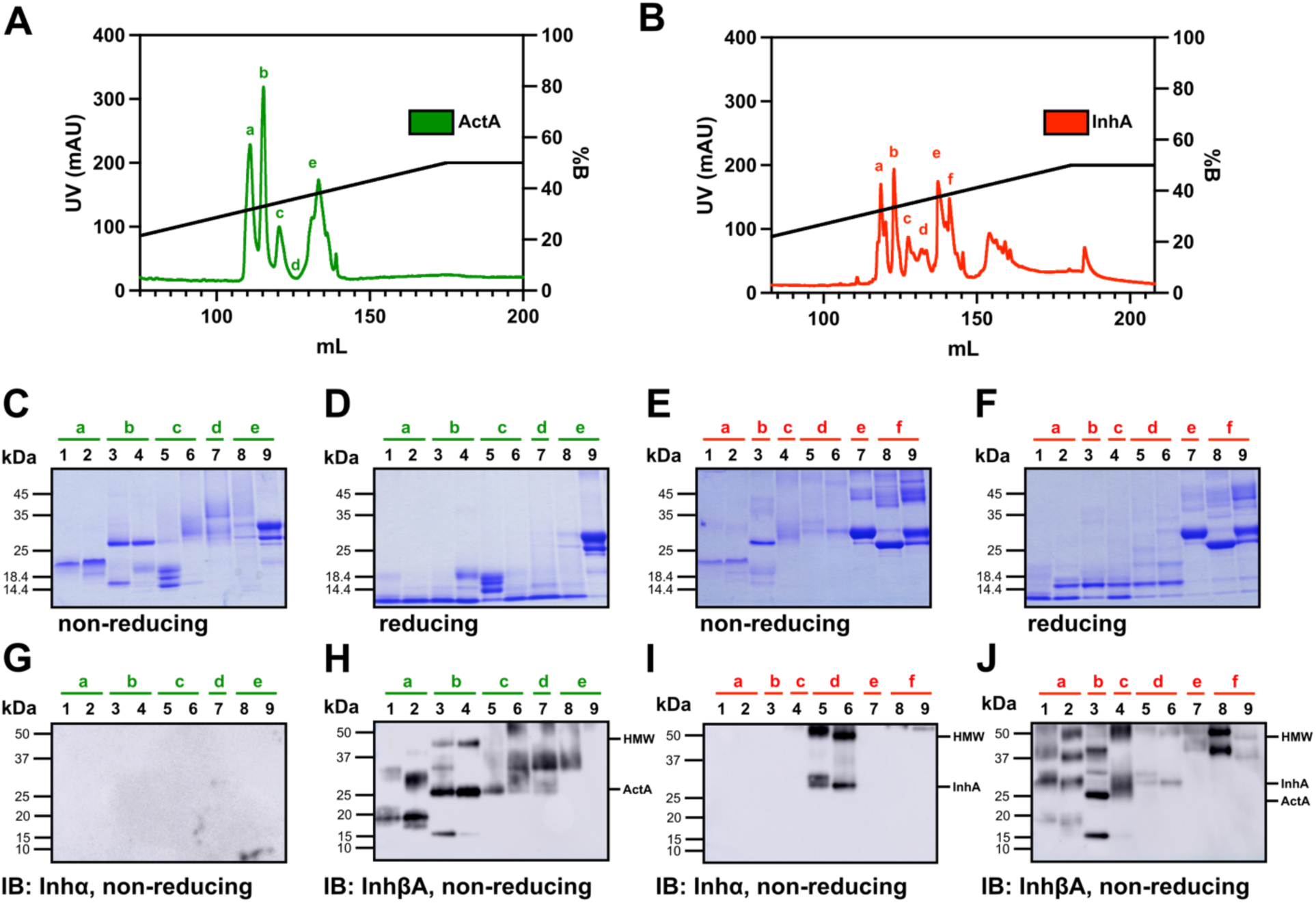
Reverse phase high-performance liquid chromatography (RP-HPLC) separates ActA from InhA. Chromatograms of RP-HPLC purifications of factor Xa-cleaved growth factor (GF) for **(A)** ActA V1 and (**B)** InhA V1, with lowercase letters corresponding to collected and resuspended fractions. SDS-PAGE gel analysis with Coomassie-staining under non-reducing and reducing conditions for fractions collected for the purifications of **(C-D)** ActA or **(E-F)** InhA (2 μg of protein/lane). Immunoblots (non-reducing) against the inhα subunit and the inhβA subunit for **(G-H)** ActA or **(I-J)** InhA fractions with a marker to denote high-molecular weight (HMW) species or growth factors (GFs) (50 ng of protein/lane).

### Purification of pro-InhA is improved by his-tagging the βA-subunit and strep-tagging the α-subunit

To reduce or eliminate co-purification of pro-ActA along with pro-InhA in the his/strep tandem purification scheme, we reasoned that placement of the strep tag on the α subunit should enable much cleaner separation of ActA and InhA, as the strep resin used here is known to exhibit much lower non-specific binding compared to Co-IMAC (34–37). Thus, we generated a tag-swapped construct with strep and his tags on the N-terminus of the pro-domain of the α and βA subunits, respectively (InhA V2 construct) (**Fig. 1D**). To evaluate the efficacy of this tagging scheme, we repeated the previous InhA purification protocol and found that pro-ActA and pro-InhA both eluted from the Co-IMAC (**Fig. 4A**) column similarly to pro-InhA V1 (compare **Fig. 4B-D, lanes 2-10** to **Fig. 2E, 2H-I**). The subsequent purification on the strep column (**Fig. 4E**) appeared to effectively separate pro-ActA from pro-InhA as the former was found only in the flow through and wash (**Fig. 4F, lanes 1-2, 7-8**) while the latter was found only in the eluate (**Fig. 4F, lanes 3, 6, 9**). The RP-HPLC purification (**Fig. 4G**) of the factor Xa-cleaved eluate from the strep column led to a larger peak corresponding to InhA (peak ‘b’, 34.5-37 %B) compared to that in the V1 purification (compare **Fig. 4H-K, peak ‘b’, lanes 2-5** to **Fig. 3B, 3I-J, peak ‘d’, lanes 5-6**). There was a marked decrease in the total amount of ActA (peak ‘a’, 33-34 % B), which we attribute to the elimination of pro-ActA with the strep purification step (**Fig. 4H-K, peak ‘a’, lane 1**).

**Figure 4:**
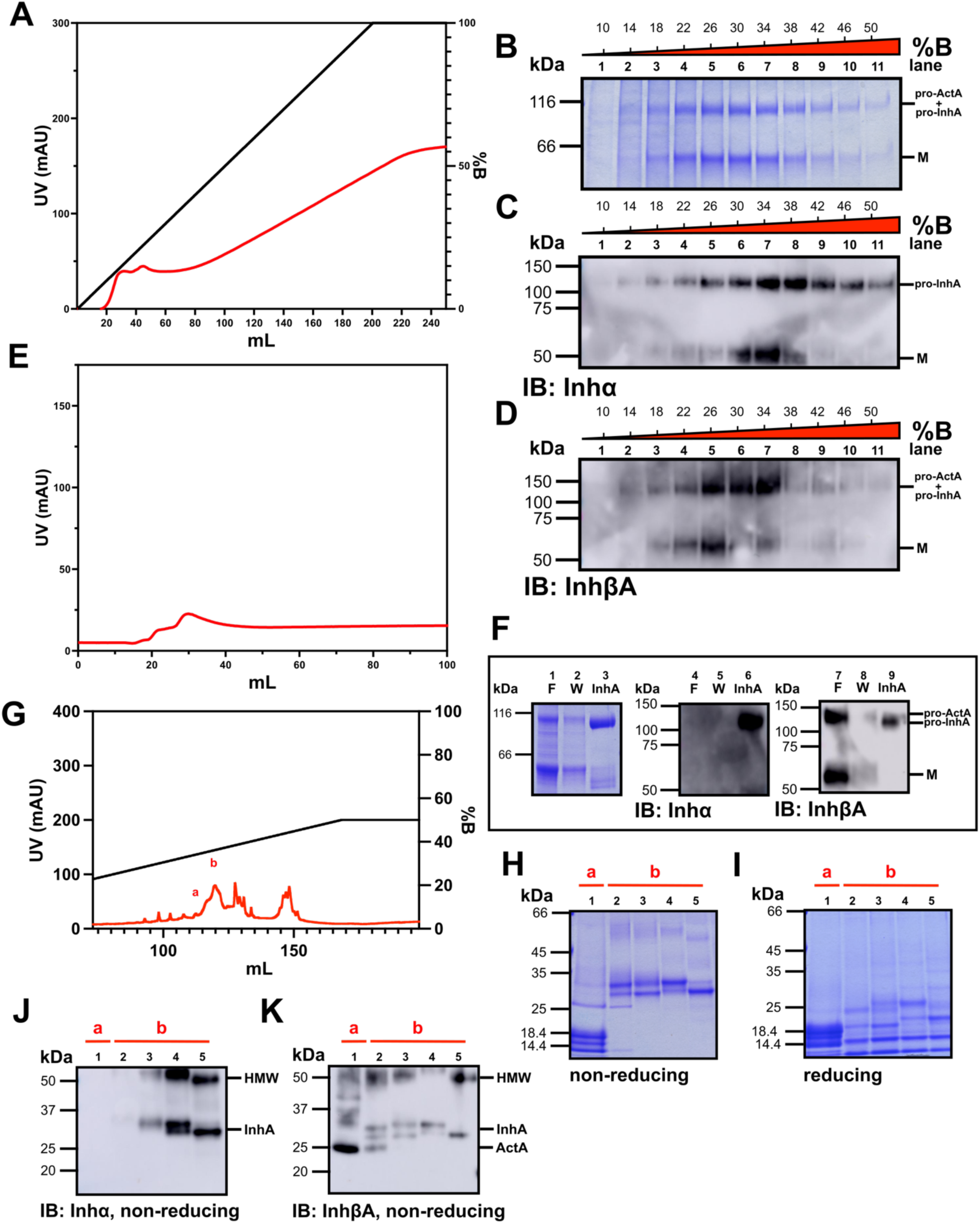
Strep-tagging the inhα subunit improves separation of ActA and InhA, prior to RP-HPLC. **(A)** Chromatogram corresponding to the Co-IMAC purification of pro-InhA V2. **(B)** Coomassie-stained SDS-PAGE gel analysis (non-reducing conditions) of eluted fractions (27.5 μL/lane) from the Co-IMAC purification, with markers for pro-ActA/pro-InhA dimers or monomer (M). Immunoblots (non-reducing) against **(C)** the inhα subunit or **(D)** the inhβA subunit of eluted fractions (0.55 μL/lane) from the Co-IMAC purification. **(E)** Chromatogram of biotin-eluted protein from the strep column. **(F)** Non-reducing SDS-PAGE gel stained with Coomassie (left) and immunoblots (middle and right) corresponding to flow-through (F), wash (W), and eluted protein from the strep affinity purification of pro-InhA. **(G)** Chromatogram of the RP-HPLC purification of InhA GF, with resuspended peaks denoted by lowercase letters. Resuspended fractions were run on SDS-PAGE gels under **(H)** non-reducing or **(I)** reducing conditions and stained with Coomassie (2 μg of protein/lane) or immunoblotted against **(J)** the inhα subunit and **(K)** the inhβA subunit, with markers to denote HMW species or GFs (50 ng of protein/lane).

### Production of wild-type InhA

We next sought to produce WT InhA from a new construct, InhA V3 (**Fig. 1E**), in which the amino acid substitutions previously employed by Goney et al. were removed (29). We followed the same tandem purification and RP-HPLC purification scheme as previously described. The protein eluted from the Co-IMAC column at 18-50 %B (**Fig. 5A-D, Supp. Fig. S1C**), as before, but there was less contaminating inhβA monomer in these fractions compared to the purification of pro-InhA V1 or V2 (compare **Fig. 5B-D, lanes 2-9** to **Fig. 2E, 2I, Fig. 4B-D**). The strep-α and his-βA tagging scheme was effective at separating pro-InhA from pro-ActA (**Fig. 5E-F**), as pro-ActA was detected in the wash and flow-through (**Fig. 5F, lanes 1-2, 7-8**) but not in the pro-InhA eluted from the strep column (**Fig. 5F, lanes 3, 6, 9**). Factor Xa-cleaved InhA GF eluted in the RP-HPLC purification (**Fig. 5G**) in peak ‘d’ (34.5-37 %B) (**Fig. 5H-K, peak ‘d’, lanes 4-7)**, with a trace amount of ActA present in peaks ‘b’ and ‘c’ (33-34 %B) (**Fig. 5H-K, peaks ‘b-c’, lanes 2-3**). The total yield from this purification was 1.14 mg of InhA per liter of medium (peak ‘d’). The purified InhA contained some contaminating HMW species, both on the Coomassie-stained SDS-PAGE gels and immunoblots (**Fig. 5H-K, lanes 4-7**). The purifications of InhA V2 and V3 (hereafter WT) were repeated a total of three times each. The average amount of WT InhA produced per liter of suspension culture volume was 1.4 mg, SD = 0.47 for V3, which was significantly greater (p = 0.0234, one-tailed t-test) than that observed for V2 (0.6 mg, SD = 0.19) (**Fig. 5L**). This suggests that the amino acid substitutions in the V2 construct reported by Goney et al. (29) were detrimental to formation of the native InhA heterodimer, at least in the expression and purification scheme that we used.

**Figure 5:**
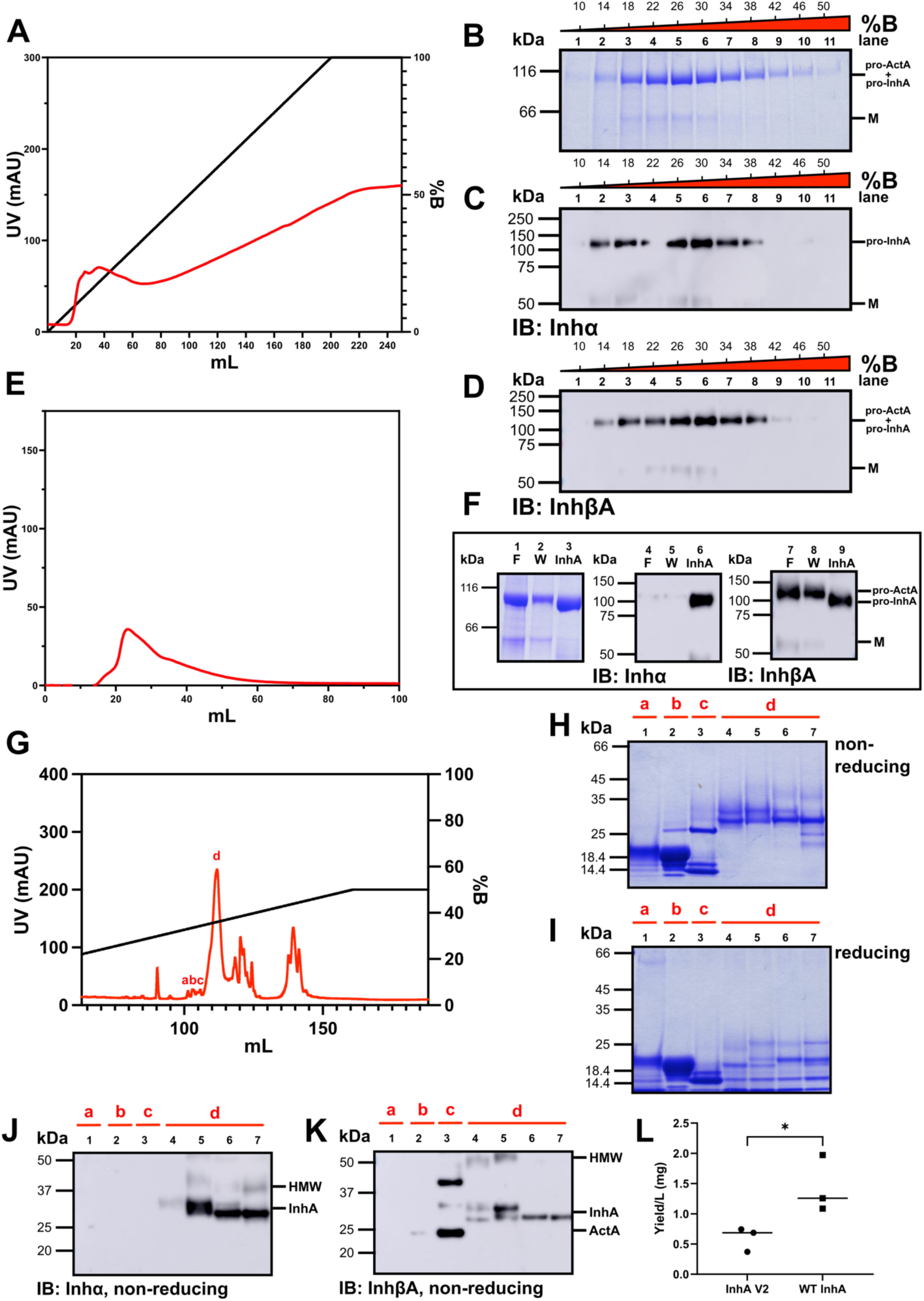
Improved InhA production with wild-type (V3) relative to V2. **(A)** Chromatogram corresponding to the Co-IMAC purification of WT pro-InhA (V3 construct). **(B)** Coomassie-stained SDS-PAGE gel analysis under non-reducing conditions of eluted protein fractions (27.5 μL/lane) from the Co-IMAC purification, with markers for pro-ActA/pro-InhA dimers and monomer (M). Purified fractions (0.55 μL/lane) were immunoblotted against **(C)** the inhα subunit and **(D)** the inhβA subunit (under non-reducing conditions). **(E)** Chromatogram of biotin-eluted protein from the strep column. **(F)** Non-reducing SDS-PAGE gel analysis with Coomassie (left) and immunoblots (middle and right) corresponding to flow-through (F), wash (W), and eluted protein from the strep-based purification of pro-InhA. **(G)** Chromatogram of the RP-HPLC purification of InhA GF, with resuspended fractions denoted by lowercase letters. Resuspended fractions were analyzed by Coomassie-stained SDS-PAGE gels under **(H)** non-reducing and **(I)** reducing conditions (2 μg of protein/lane). Fractions were immunoblotted (non-reducing) against **(J)** the inhα subunit and **(K)** the inhβA subunit (50 ng of protein/lane). **(L)** Purifications of mutant InhA V2 and WT InhA were repeated, and total InhA yields were compared (n = 3, each point is an experimental replicate). For **(L)** and Fig. 6L, graphed data are presented as individual values with a bar depicting the mean and analyzed by one-tailed unpaired Student’s t-test. (* P < 0.05).

### InhB can be produced and purified using a similar tagging and purification scheme, but with lower yield than InhA

Owing to our success producing large amounts of WT InhA, we adapted the same approach to produce WT InhB (**Fig. 1F**), though on a larger scale than InhA in anticipation of a lower overall yield (informed by our own unpublished observations^1^). Pro-InhB was eluted from the Co-IMAC column (**Fig. 6A**) at a similar imidazole concentration (14-50 %B) as observed for pro-InhA (compare **Fig. 6B-D, lanes 2-10** to **Fig. 5B-D**). The strep column purification (**Fig. 6E**) successfully separated pro-ActB (**Fig. 6F, lanes 1-2, 7-8**) from pro-InhB (**lanes 3, 6, 9**). We observed trace amounts of pro-InhB detectable by probing against inhα that were not captured by the strep column, although this was not reproducible (**Fig. 6F, lanes 4-5**). Factor Xa-cleaved InhB GF eluted from the RP-HPLC column (**Fig. 6G**) at a higher concentration of acetonitrile (peak ‘a’, 36-39 %B) than observed for InhA (peak ‘d’ 34.5-37 %B) (compare **Fig. 6G-K, peak ‘a’, lanes 1-3** to **Fig. 5G-K, peak ‘d’, lanes 4-7**). No contaminating ActB GF was observed in the purified fractions (**Fig. 6H-K, lanes 1-3**). We observed both glycoforms of mature InhB, as well as a band at ∼25 kDa that was detected by the anti-inhα antibody, but not by the anti-inhβB antibody (**Fig. 6J-K, lanes 1-3**). There were also contaminating HMW species detected by antibodies against both subunits (**Fig. 6J-K, lanes 1-3**). The average yield of purified InhB was 0.093 mg per liter of suspension culture (n = 3, SD = 0.043), more than 10-fold lower than the average yield for InhA (p = 0.0039, one-tailed t-test) (**Fig. 6L**).

**Figure 6:**
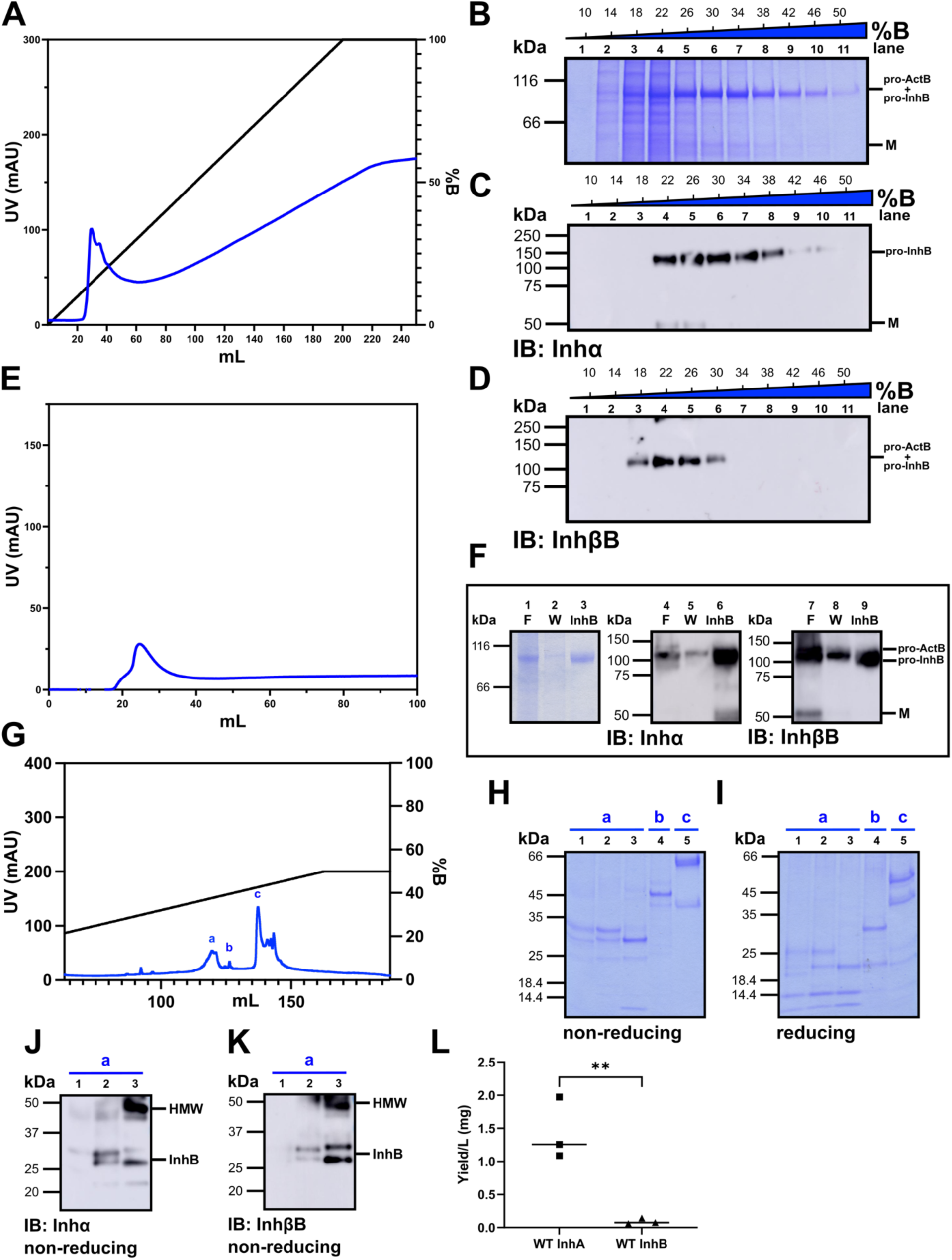
InhB is produced with lower efficiency than InhA. **(A)** Chromatogram corresponding to the Co-IMAC purification of pro-InhB. **(B)** Coomassie-stained SDS-PAGE gel analysis (non-reducing) of eluted protein fractions (27.5 μL/lane) from Co-IMAC purification and immunoblots (0.55 μL/lane) (non-reducing) against **(C)** the inhα subunit or **(D)** the inhβB subunit. **(E)** Chromatogram of biotin-eluted protein from the strep affinity column. **(F)** The corresponding non-reducing Coomassie-stained SDS-PAGE gel (left) and immunoblots (middle and right) corresponding to flow-through (F), wash (W), and eluted protein from the strep purification of pro-InhB. **(G)** Chromatogram corresponding to the RP-HPLC purification of InhB GF, with resuspended fractions denoted by lowercase letters. Resuspended fractions were analyzed by SDS-PAGE gels with Coomassie staining under **(H)** non-reducing and **(I)** reducing conditions (1 μg of protein/lane). Fractions were immunoblotted (non-reducing) against **(J)** the inhα subunit and **(K)** the inhβB subunit, with markers to denote HMW species or GF (100 ng of protein/lane). **(L)** Total InhA and InhB yields were compared (n = 3, each point is an experimental replicate). (** P < 0.01).

### InhA and InhB purified by our optimized procedure function similarly to commercially available inhibins

We next compared the purity and bioactivity of our WT InhA and InhB preparations to commercially available human inhibins. We first tested 10 ng (as determined by UV absorbance using the calculated extinction coefficient) of our WT human InhA and a sample of carrier-free human InhA from R&D Systems (**Fig. 7A-B**). We again observed our WT InhA was composed of both the 31- and 34-kDa glycoforms (**Fig. 7A-B, lane 2**), whereas the R&D InhA consisted predominantly of the 31-kDa form (**Fig. 7A-B, lane 1**) (2, 38). We observed slightly stronger signal across glycoforms for our InhA compared to R&D InhA, likely explained by inaccuracies in the concentration due to contaminants in both preparations (**Fig. 7A-B**).

**Figure 7:**
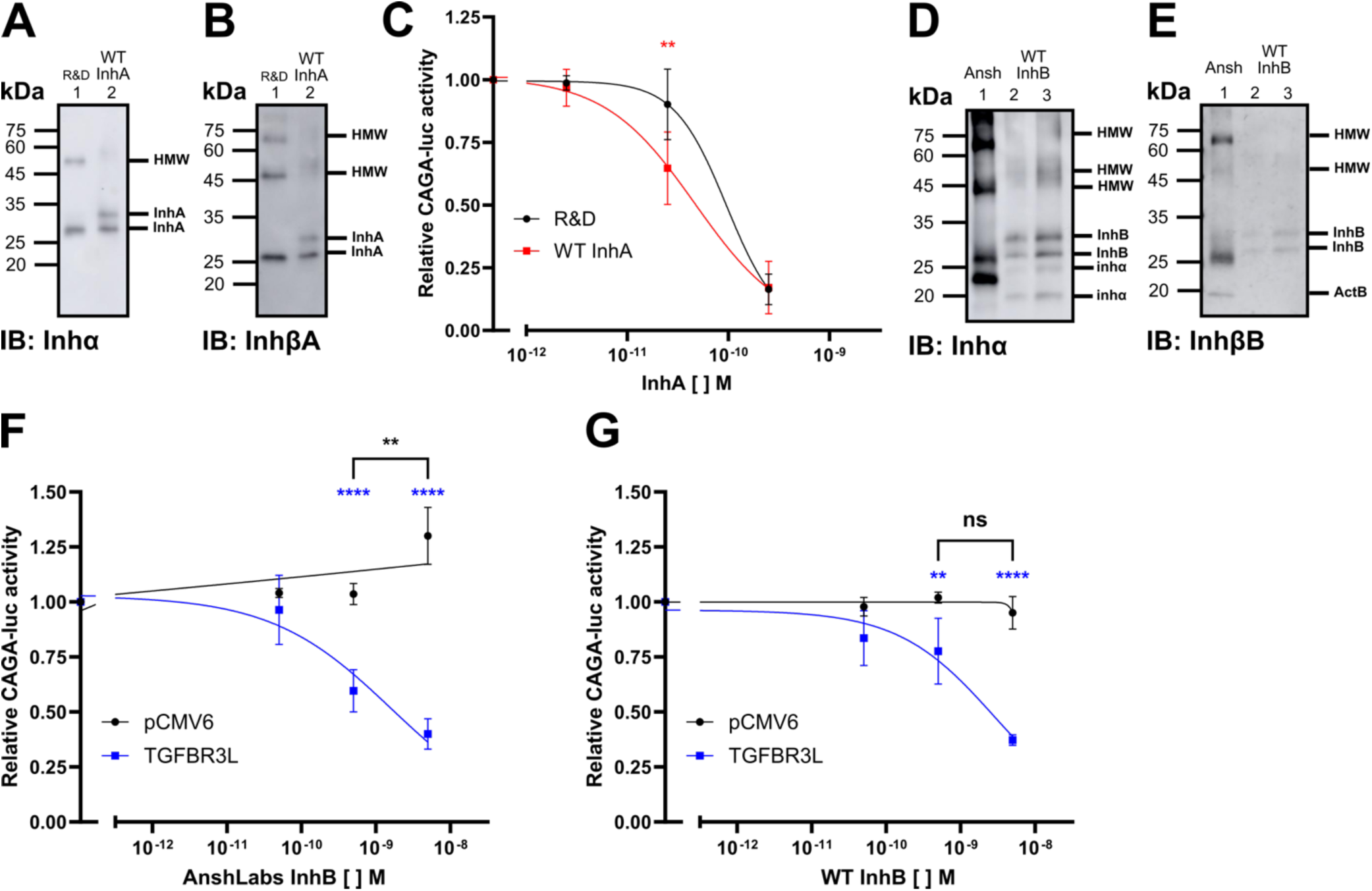
Purified inhibin A and B preparations antagonize ActA as effectively as industry standards. Equal amounts of total protein (10 ng/lane) of human R&D Systems InhA or WT InhA (produced from the V3 construct) were immunoblotted (non-reducing) against **(A)** the inhα subunit and **(B)** the inhβA subunit. **(C)** CHO cells transiently transfected with rat betaglycan ZPC and the CAGA-luc promoter-reporter were treated with 1 nM ActA (R&D) and increasing concentrations of R&D Systems InhA or WT InhA. Reporter activity was normalized to the ActA response, in the absence of InhA (n = 4 experimental replicates). AnshLabs human InhB (30 ng, lane 1), and 30 or 60 ng of InhB (lanes 2-3) were immunoblotted (non-reducing) against **(D)** the inhα subunit and **(E)** the inhβB subunit. InhB activity was assessed in CHO cells transfected with empty vector or murine TGFBR3L and the CAGA-luc promoter reporter. Cells were treated with 1 nM ActA (R&D) alone, or in the presence of the indicated amounts of **(F)** AnshLabs human InhB or **(G)** purified InhB. Reporter activity was normalized to the ActA response, in the absence of InhB (n = 3 or 4 experimental replicates). For **(C)**, **(F)**, and **(G),** data represent mean values (error bars depicting SD), and were analyzed by two-way ANOVA, followed by Sidak’s multiple comparisons test (not significant (ns), * P < 0.05, ** P < 0.01, **** P < 0.0001).

To test the activity of the inhibins, we adjusted the concentration to reflect the amount of mature GF, calculated using a standard curve calculated from SEC purified InhA (**Supp. Fig. S2A-D**). We transfected Chinese hamster ovary (CHO) cells with the SMAD3/4 responsive CAGA-luc promoter-reporter and either betaglycan ZPC or empty vector. Betaglycan increases the sensitivity of these cells to inhibins (23). Then, we treated cells with 1 nM ActA (R&D Systems) and increasing concentrations of InhA. In the presence of betaglycan ZPC, our non-SEC purified WT InhA antagonized ActA similarly to R&D human InhA, although ours displayed greater inhibition at 0.025 nM (**Fig. 7C**, **Table 1**). No ActA activity was observed in either InhA preparation (**Supp. Fig. S2E-F**).

**Table 1.**
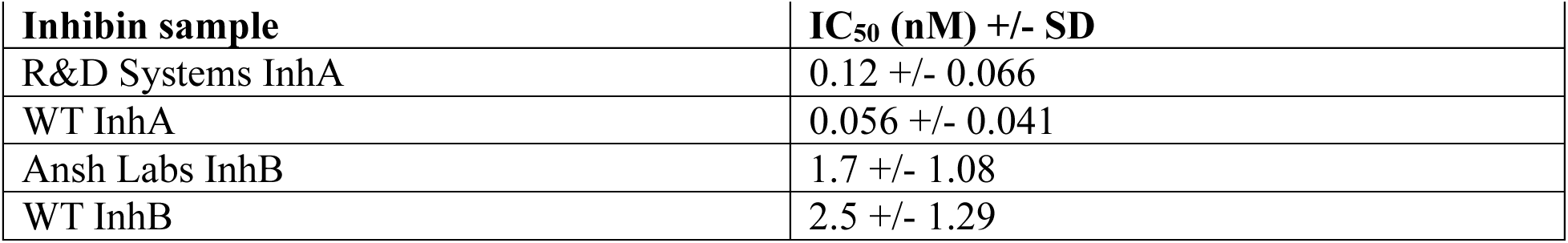
IC_50_s for wild-type inhibin A and B and their commercial sources.

| **Inhibin sample** | **IC<sub>50</sub> (nM) +/- SD** |
| --- | --- |
| R&D Systems InhA | 0.12 +/- 0.066 |
| WT InhA | 0.056 +/- 0.041 |
| Ansh Labs InhB | 1.7 +/- 1.08 |
| WT InhB | 2.5 +/- 1.29 |

We compared our InhB to commercially available human InhB from AnshLabs (**Fig. 7D-G**). No ActB was detected in our InhB preparation (**Fig. 7E, lanes 2-3)**, unlike the AnshLabs InhB **(Fig. 7E, lane 1)**. After correcting the concentration to reflect the amount of mature GF in the two preparations, both InhBs antagonized ActA stimulation of CAGA-luc with comparable potency in the presence of co-transfected inhibin B co-receptor, TGFBR3L, in CHO cells (**Fig. 7F-G**, **Table 1, Supp. Fig. S3A-D**). In the absence of TGFBR3L, we observed no ActB activity in our InhB. In contrast, AnshLabs InhB stimulated CAGA-luc at the highest concentration tested (**Fig. 7F-G**).

## Discussion

We developed a new and effective method of purifying inhibins for functional and biophysical studies. Our protocol does not use any proprietary reagents or cell lines and can be used to generate milligram-scale quantities of highly purified WT InhA, providing greater flexibility than other established protocols (29, 31, 32).

### Effective and rational use of the dual tagging scheme is essential to purify InhA from ActA

Appropriate use of the Co-IMAC column and the dual tagging scheme (strep-tagged α and his-tagged β) was essential to purify InhA away from ActA. The Co-IMAC column was ineffective at separating InhA from ActA when used with his-tagged inhα because inhβA lacking a his-tag bound non-specifically to the column almost as well as inhα harboring a his-tag. This non-specific binding was not previously reported in (29) and may further restrict the applicability of that protocol. Since activin contamination was not assessed via Coomassie-stained SDS-PAGE gels or immunoblots, it is unknown whether the inhibin purified from CM in (29) was free of contaminating activin. We reasoned that replacing the his-tag on the α-subunit with a strep tag, which is known to bind with high specificity to strep columns (39–41), would eliminate the vast majority of contaminating ActA in our InhA preparations. This proved to be effective with our InhA V2 and V3 constructs, which drastically reduced the amount of ActA eluting from the RP-HPLC column relative to that observed for InhA V1.

### InhA is produced with higher yield without inhα^S334I^, inhβA^A347H^, inhβA^M334A^, and the α-subunit ‘SCUT’ site modifications

We successfully produced InhA with previously described substitutions (inhα^S334I^, inhβA^A347H^, inhβA^M334A^, and the α-subunit ‘SCUT’ site) (29, 30), but found that the yields of the InhA GF were improved 2.5-fold when excluding these modifications. This may be due to elimination of inhβA^A347H^, as the A347H substitution increased the amount of inhβA monomer with V1 and V2 of pro-InhA compared to V3, which lacks this substitution. Thus, in addition to impairing inhβA homodimerization, as previously shown (29, 30), inhβA^A347H^ might also impair inhβA-inhα heterodimerization, accounting for the reduced yield. It is possible that the other included substitutions had a negative impact on InhA production; however, we did not individually assess them. Hence, further studies are required to determine whether this residue, centrally located in the dimer interface, is solely responsible. Additionally, the prior study relied on endogenous furin processing (29), while we used orthogonal factor Xa to cleave the pro-domain. Thus, we cannot rule out the possibility that the substitutions behave differently in the endogenous context. However, successful production of this modified InhA demonstrated that this system could be used to generate variant InhA for structural and biophysical studies.

### InhB production is feasible, but reduced compared to InhA

We found that our system could be adapted for producing InhB without the substitutions suggested by Goney et al. (29, 30). We were therefore able to compare the relative efficiency of producing InhB to InhA, which to our knowledge has not previously been reported. We observed that the overall yield of InhB was reduced by at least 10-fold compared to InhA. One interpretation is that InhB has a poorer propensity to fold. In support of this, we find AlphaFold3 model confidence decreases when pro-inhβB is present in homo- and hetero-dimers in place of pro-inhβA (**Supp. Fig. S4**) (42). Perhaps the inhβB pro-domain does not promote effective heterodimerization. This could explain the presence of ∼25 and ∼20 kDa bands detected by the anti-inhα antibody, which might arise from the inability of inhβB to fold and form a covalent heterodimer with inhα (**Fig. 6J, lanes 1-3, Fig. 7D, lanes 2-3**). These bands have been reported before and shown to correspond to di- and mono-glycosylated inhα monomer, consistent with our immunoblotting results (10). If pro-inhβB has a poor propensity to fold and form a covalent heterodimer with pro-inhα, we may be able to modify and optimize its folding (e.g., swapping in regions of pro-InhβA). If residues within the GF-GF interface impair folding and heterodimerization, we may be limited in which substitutions can be made to increase production of proteins involving this subunit while maintaining near WT sequence.

### Limitations of our system, future improvements, and applications

One limitation of the inhibin production system presented here is that the inhibins produced are not strictly WT. We substituted G316 close to the N-terminus of the mature βA-subunit with lysine to eliminate a potential factor Xa secondary cleavage site, Gly-Lys, and we made similar modifications G298A and R299K to the βB subunit (to eliminate a Gly-Arg secondary site) (**Fig. 1E-F**). Despite these substitutions, the InhA and InhB we produced displayed comparable bioactivity to commercially available InhA and InhB. The current system might be improved by replacement of the factor Xa processing site with the processing site of another protease, such as tobacco etch virus protease or human rhinovirus 3C protease, which have greater specificity, obviating the need for the βA G316K substitution (43, 44). It remains to be determined whether the βB G298A/R299K modifications or absence of endogenous furin processing impaired InhB production, or whether InhB production can be improved by optimizing either the plasmid or the transfection method. Future studies will assess production using the Expi293™ PRO Expression System, either by transient transfection or viral infection.

While our V2 (and V3) tagging and tandem purification scheme was highly effective in eliminating contaminating activins, we detected trace amounts. These could be eliminated using RP-HPLC. We also observed contaminating HMW species in fractions corresponding to the factor Xa-cleaved mature GFs. This was more apparent by immunoblotting than Coomassie staining. Despite this, our inhibins exhibited similar bioactivity to commercially available standards (which appeared similarly affected by contaminating species). Hence, we think that our method is sufficient for producing quality InhA in quantities required for both functional (in cells and animals) and biophysical studies. The InhB produced is of similar quality, but refinements may be needed to improve yields. We expect our bis-cistronic expression and purification system will work for production and purification of other heterodimeric TGF-β family ligands, such as BMP9/10, BMP2/6, BMP2/7, BMP4/7, Vg1/Nodal, ActAB, and TGF-β1/2 (45–50). Producing pure heterodimers for use in cell-signaling and biophysical assays will allow us to assess the function of TGF-β family heterodimers in physiology (2, 16, 45–47, 49, 51–53).

### Experimental Procedures

The key resources used for this study are summarized in Table 2.

**Table 2.** Key resources.

| **Name** | **Details** |
| --- | --- |
| Infusion SNAP assembly mix | Cat# 638948, Takara Bio, USA |
| 400 mL Amicon stirred cell | Cat# UFSC40001, EMD Millipore, Burlington, MA, USA |
| HisPur NTA agarose | Cat# 88222, ThermoFisher Scientific, Waltham, Massachusetts, USA |
| CoCl<sub>2</sub> | Cat# C8661-100G, Sigma, Darmstadt, Germany |
| ÄKTAprime plus | Cytiva, Piscataway, NJ, USA |
| Strep-Tactin XT 4Flow high-capacity resin | Cat# 2-5030-010, IBA, Göttingen, Germany |
| D-biotin | Cat# B-950-25, GoldBio, St.-Louis, Missouri, USA |
| Bovine factor Xa | Cat# BFXa, Enzyme Research Laboratories, South Bend, Indiana, USA |
| polyvinylidene difluoride (PVDF) 0.1 uM filter | Cat# UFC30VV00, Ultrafree, Millipore, Darmstadt, Germany |
| C4 RP-HPLC column (4.6 mm x 250 mm) | Cat# 214TP104, Grace/Vydac, Hesperia, CA, USA |
| ÄKTA pure | Cytiva, Piscataway, NJ, USA |
| Nanodrop 2000 | ThermoFisher Scientific, Waltham, Massachusetts, USA |
| Superdex 75 increase 10/300 size-exclusion chromatography column (SEC) | Cat# 17517401, Cytiva, Piscataway, NJ |
| SurePAGE™, Bis-Tris, 10x8 resolving gel | Cat# M00659, GenScript, Piscataway, NJ, USA |
| Tris-MOPS-SDS running buffer | Cat# M00138, GenScript, Piscataway, NJ, USA |
| Epson Perfection 1650 | Epson, Los Alamitos, California, USA |
| PVDF transfer membranes | Cat# 88520, Thermo Fisher Scientific, Waltham, Massachusetts, USA |
| Anti-inhibin α (R1) | Cat# ab14087, RRID:AB_2811124, Abcam, Cambridge, UK |
| Monoclonal mouse anti-inhibin βA | Cat# sc-166503, RRID:AB_2125854, lot# C1020, Dallas, TX, USA |
| Monoclonal mouse anti-inhibin βB | Cat# sc-376971, RRID:AB_3677370, lot# D0219, Dallas, TX, USA |
| Horseradish peroxidase–conjugated anti-mouse goat anti-mouse | Cat# ab205719, RRID:AB_2755049, Abcam, Cambridge, UK |
| Amersham enhanced chemiluminescence Prime | Cat# RPN2232, Cytiva, Piscataway, NJ |
| Amersham Imager 600 | GE Healthcare, Chicago, IL, USA |
| Unstained protein molecular weight marker | Cat# 26610, ThermoFisher Scientific, Waltham, Massachusetts, USA |
| Precision Plus Dual Color Standards | Cat# 161-0374, Bio-Rad, Hercules, California, USA |
| Protran nitrocellulose membranes | Cat# GE 10600001, MilliporeSigma, Burlington, Massachusetts, USA |
| Monoclonal mouse anti-inhibin βA | Cat# MAB-3381, RRID:AB_2125864, R&D systems, Minneapolis, MN, USA |
| Monoclonal mouse anti-inhibin βB | Cat# AB-306-AA042; RRID:AB_2756475, AnshLabs, Webster, TX, USA |
| HRP-conjugated secondary antibody goat anti-mouse | Cat# 1706516, RRID:AB11125547, Bio-Rad Laboratories, Saint-Laurent, QC, Canada |
| Activin A | Cat# 338-AC, R&D systems, Minneapolis, MN, USA |
| Human InhA | Cat# 8506-AB-010/CF, R&D systems, Minneapolis, MN, USA |
| Human InhB | Cat# AG-306-AI100, AnshLabs, Webster, TX, USA |

### DNA constructs

The N-terminal myc-tagged rat betaglycan ZPC was amplified from synthetic double-stranded DNA fragments (Twist Bioscience, San Francisco, CA, USA) using primers that inserted a 5′ *EcoR*I site and a 3’ *Hind*III site. The fragment encoded the rat betaglycan signal peptide (M1-R30) followed by a myc tag and the coding sequence of the ZPC, transmembrane domain, and cytoplasmic tail (G590-A853, Uniprot P26342). The amplicon was inserted into the pCMV5 backbone.

The coding sequences for human inhβA (UniProt P08476), inhβB (UniProt P09529), and inhα (UniProt P05111) with the features for the V1, V2, and V3 constructs shown in **Fig. 1**, but with an added Kozak sequence on the 5’ end and the native signal peptide replaced with the rat serum albumin signal peptide, were synthesized as double-stranded DNA fragments (Twist Bioscience). These fragments were inserted between the 5′ *EcoR*I and 3′ *Xba*I restriction sites downstream of the CMV promoter in plasmid pEG (33). The bis-cistronic plasmids encoding V1, V2, V3 human InhA and V3 InhB were generated by amplifying the pEG inhα plasmid with primers, 5’-GAATCGATACTAGTATACAAACAAAGCTGGCTATGGCAG-3’ and 5’-TGCATCAGCTGCTAGCGCTCAAGCAGTGATCAGATCCAGACATG-3’, which anneal to pEG vector sequences flanking the coding cassette. The resulting PCR product, 3.8 kb in size, was inserted into the pEG-inhβA or pEG-inhβB plasmid digested with *Nhe*I and *BstZ*17I using In-Fusion SNAP Assembly mix (Takara Bio). The sequences of all plasmids were verified by nanopore-based whole plasmid sequencing (Plasmidsaurus).

### Protein expression and tandem affinity purification

Protein expression in Expi293 suspension-cultured HEK293 (RRID:CVCL_D615) cells was carried out, unless otherwise specified, on a scale of 300 mL as previously described (54). Conditioned media containing the expressed protein was collected 72 h post-transfection, supplemented with protease inhibitors (0.1 mM PMSF, 1 mM benzamidine, 1 μM pepstatin, and 1 μM leupeptin), concentrated to 120 mL using a 400 mL Amicon stirred cell (Millipore), and dialyzed at 4°C overnight against 4 L of buffer containing 50 mM sodium phosphate and 150 mM NaCl, pH 8.0 with protease inhibitors. On the following day, the dialyzed conditioned media was loaded onto a 20 mL immobilized metal affinity column [HisPur NTA agarose (Thermo Fisher Scientific) charged with CoCl2 (Sigma) (Co-IMAC column)] on an ÄKTAprime plus (Cytiva) and then washed with buffer until the UV absorbance at 280 nm returned to baseline. The combined flow-through and wash were collected, if necessary. Protein was eluted from the column with a 0-0.5 M imidazole gradient (pH 8.0) over 200 mL and collected in 8 mL fractions. Chromatograms were aligned to the start of the gradient. The combined flow-through and wash from the Co-IMAC column were concentrated to 80 mL, approximating the volume of protein-containing fractions.

If the protein contained a strep tag, protein-containing fractions were combined and dialyzed overnight against 4 L of strep buffer containing 50 mM Tris, 150 mM NaCl, 1 mM EDTA, pH 8.6 with protease inhibitors. The next day, protein was loaded onto an equilibrated Strep-Tactin XT column (IBA Lifesciences) on an ÄKTAprime plus. Flow-through was collected and re-loaded onto the column three times, before a 250 mL wash with strep buffer. Collected flow-through and wash were concentrated to 1 mL, if needed, to compare the amount of unbound protein to the eluate. Bound protein was eluted with 50 mM D-biotin (GoldBio) in strep buffer. Chromatograms were aligned to the start of the elution. 5 mL fractions were collected, and protein-containing fractions were combined and concentrated to 1 mL and dialyzed three times for 12 h against 4 L of buffer containing 20 mM Tris, 50 mM NaCl, and 1 mM CaCl_2_, pH 6.5. The tandem purified pro-inhibin was cleaved for 16 h at 25 °C with bovine factor Xa (Enzyme Research Laboratories), 10 μg for each 1 mg of protein.

Following factor Xa cleavage, samples were brought to pH 4.0 with glacial acetic acid and then filtered through a 0.1-μm polyvinylidene difluoride (PVDF) filter (Millipore) to remove precipitate. Protein was loaded onto a C4 RP-HPLC column (Grace/Vydac, 4.6 mm x 250 mm) on an ÄKTA pure 25M (Cytiva) equilibrated in buffer A (94.9% H_2_O, 5% acetonitrile, 0.1% trifluoroacetic acid), and eluted with a linear gradient from 10-50% buffer B (99.9% acetonitrile, 0.1% trifluoroacetic acid) over 140 mL and collected in 2 mL fractions. Fractions were frozen at -80° C and lyophilized overnight. Lyophilized fractions were resuspended in 4 mM HCl. Protein concentrations were determined by measuring the absorbance at 280 nm (NanoDrop 2000, Thermo Fisher Scientific) and the calculated extinction coefficient.

To further purify InhA, 50 μg of RP-HPLC purified inhibin A was loaded onto an equilibrated Superdex 75 increase 10/300 size-exclusion chromatography (SEC) column (Cytiva) on an ÄKTA pure 25L (Cytiva) and 0.5 mL fractions were collected. The buffer was 1 mM HCl and 200 mM NaCl (pH 3.0). Fractions corresponding to inhibin were pooled and the buffer was adjusted to 4 mM HCl, then the volume was brought to 100 μL. Protein concentration was determined by measuring the UV absorbance at 280 nm, as described above.

### SDS-PAGE and immunoblots used during protein purification

Two times concentrated tricine-SDS sample buffer [30% 3.0 M Tris-HCl (pH 8.45), 24% glycerol, 8% SDS (w/v), 0.009% Coomassie Blue G, 0.005% Phenol Red] was added to protein samples, which were then resolved by SDS-PAGE on an 8-16% SurePAGE™, Bis-Tris, 10x8 resolving gel (GenScript) in Tris-MOPS-SDS running buffer (GenScript). For reducing gels, 100 mM DTT was added to the sample buffer. Gels subject to staining were placed in stain (45% methanol, 45% H_2_O, 10% acetic acid, and 0.25% (w/v) Coomassie Blue R-250) for 30 min with agitation, and destained overnight (45% methanol, 45% di-H_2_O, 10% acetic acid). Following destaining, gels were soaked in drying solution (5% glycerol/30% methanol), dried in cellophane, and scanned the following day (Epson Perfection 1650, Epson).

For immunoblotting, proteins were transferred to PVDF transfer membranes (Thermo Fisher Scientific) in transfer buffer (25 mM tris, 192 mM glycine (pH 8.3), 20% methanol, and 0.01% SDS), blocked with 5% bovine serum albumin (BSA) (w/v) in Tris-buffered saline (TBS) (50 mM NaCl and 10 mM Tris (pH 7.6)) containing 0.01% (v/v) Tween-20 (TBST) and incubated for 1 h with agitation with the indicated primary antibodies diluted in blocking buffer. The monoclonal mouse anti-inhibin α (R1) (1:200) was purchased from Abcam (RRID:AB_2811124). The monoclonal mouse anti-inhibin βA (RRID:AB_2125854) (1:1000) and anti-inhibin βB (RRID:AB_3677370) (1:500) were purchased from Santa Cruz biotechnology. Then, membranes were washed in TBST and incubated in horseradish peroxidase–conjugated anti-mouse (1:5000, goat anti-mouse (Abcam, RRID:AB_2755049)) diluted in TBST for 1 h at room temperature with agitation. Membranes were washed in TBST, and bands were visualized with Amersham enhanced chemiluminescence Prime (Cytiva) on an Amersham Imager 600 (GE Healthcare).

Unstained protein molecular weight marker (Thermo Fisher Scientific) was used for Coomassie-stained SDS-PAGE gels. Precision Plus Dual Color Standards (Bio-Rad) were used for immunoblots.

### Immunoblotting of inhibins in preparation for cell activity assays

To quantify or visualize the amount of mature ligand in inhibin preparations by immunoblot for **Fig. 7** and **Supp. Fig. S2-3**, 10 to 150 ng of purified inhibin were diluted in five times concentrated Laemmli buffer (250 mM Tris (pH 6.8), 10% SDS, 50% glycerol, and 0.2% bromophenol blue, pH 6.8) and resolved on a 12% resolving gel (29:1 acrylamide/bis-acrylamide in running buffer, 25 mM Tris, 250 mM glycine, and 0.1% SDS, pH 8.3). Then, proteins were transferred to Protran nitrocellulose membranes (MilliporeSigma) in Towbin buffer (25 mM Tris, 192 mM glycine, 20% methanol, pH 8.3), blocked with 5% bovine serum albumin (w/v) in TBST, and incubated in primary antibody, either the R1 antibody described earlier, the monoclonal mouse anti-inhibin βA (R&D Systems, RRID:AB_2125864, 1:1000), or the anti-inhibin βB (AnshLabs, RRID:AB_2756475, 1:1000) diluted in blocking buffer, for 1 h at room temperature. Then, membranes were washed twice in TBST and incubated for 1 h at room temperature in HRP-conjugated goat anti-mouse secondary antibody (1:5000, Bio-Rad). Following two more washes in TBST, membranes were soaked in enhanced chemiluminescence substrate (PerkinElmer), and bands were imaged using an Amersham Imager 600 (GE Healthcare). In some cases, immunoblots were stripped with 0.2 M NaOH (2 x 20 min wash) followed by a 20-min wash in TBST. Membranes were then probed again (always R1 first, then anti-inhβA or βB).

### Quantification of mature GF

UV approaches were not an accurate way to determine the concentration of mature GF in inhibin samples because there were contaminating species observed in all inhibin preparations. The amount of mature GF in inhibin preparations was quantified by comparing the aggregate band intensity of both glycoforms to a standard curve of the SEC purified InhA material blotted with the α-subunit antibody (**Supp. Fig. S2C-D, S3A-D**). Band intensities were quantified using FIJI software (55).

### Cell activity assays

Chinese hamster ovary (CHO) (RRID:CVCL_0213) cells were seeded at a density of 30,000 cells/well in 48-well plates (23). The next day, cells were transfected with 200 ng of CAGA-luc, and either 10 or 100 ng of empty vector or expression vector for betaglycan ZPC or murine TGFBR3L (23), with a 1:3 ratio of PEI. Cells were serum-starved 24 h after transfection, for 24 h. For InhA activity assays: following serum-starvation, cells were treated for 6 h with vehicle, 1 nM of ActA (R&D Systems), and equal concentrations of mature growth factor for human InhA (R&D Systems) or WT InhA. To compare InhA from R&D and our WT InhA, we assayed concentrations: 0.0025, 0.025, and 0.25 nM. For InhB: after serum-starvation, cells were treated for 6 hours with vehicle, 1 nM of ActA (R&D Systems), and equal concentrations of mature growth factor for human InhB (AnshLabs) or WT InhB [0.05, 0.5, or 5 nM]. Then, luciferase activity was assayed as previously described (23). Conditions were performed in technical duplicates, and experimental triplicates or quadruplicates.

### Statistical analysis

Inhibin yields were compared by one-tailed unpaired Student’s t-test. CHO cell luciferase assay results were analyzed by two-way analysis of variance (ANOVA), followed by Sidak’s multiple comparisons test. Statistical analysis was completed using GraphPad Prism 10 software. P < 0.05 was considered statistically significant.

### AlphaFold3 predictions

AlphaFold3 modelling was done using the endogenous amino acid sequences for immature and mature activins and inhibins (2, 42). Predictions were repeated three times and model confidence scores were calculated and averaged.

## Data Availability

All data supporting the findings of this study are included in the article and its Supporting Information.

## Supporting Information

This article contains supporting information.

## Acknowledgements

We would like to thank P. Morris (The Population Council) for CHO cells. We would like to thank A.N. Bayne, N. Croteau, and J.F. Trempe (McGill University) for helping with the SEC purification of InhA.

## Footnotes

1 Buddle, E.R.S. Hinck, C.S., and Hinck. A.P. (2024).

## References

1. Bernard DJ, Fortin J, Wang Y, Lamba P. Mechanisms of FSH synthesis: what we know, what we don’t, and why you should care. Fertil Steril. 2010;93(8):2465–85.

2. Makanji Y, Zhu J, Mishra R, Holmquist C, Wong WP, Schwartz NB, et al. Inhibin at 90: from discovery to clinical application, a historical review. Endocr Rev. 2014;35(5):747–94.

3. Matsuo H, Baba Y, Nair RM, Arimura A, Schally AV. Structure of the porcine LH- and FSH-releasing hormone. I. The proposed amino acid sequence. Biochem Biophys Res Commun. 1971;43(6):1334–9.

4. Schally AV, Arimura A, Baba Y, Nair RM, Matsuo H, Redding TW, et al. Isolation and properties of the FSH and LH-releasing hormone. Biochem Biophys Res Commun. 1971;43(2):393–9.

5. Ongaro L, Zhou X, Wang Y, Schultz H, Zhou Z, Buddle ERS, et al. Muscle-derived myostatin is a major endocrine driver of follicle-stimulating hormone synthesis. Science. 2025;387(6731):329–36.

6. Wildt L, Hausler A, Marshall G, Hutchison JS, Plant TM, Belchetz PE, et al. Frequency and amplitude of gonadotropin-releasing hormone stimulation and gonadotropin secretion in the rhesus monkey. Endocrinology. 1981;109(2):376–85.

7. Kumar TR, Wang Y, Lu N, Matzuk MM. Follicle stimulating hormone is required for ovarian follicle maturation but not male fertility. Nat Genet. 1997;15(2):201–4.

8. Sengle G, Ono RN, Sasaki T, Sakai LY. Prodomains of transforming growth factor beta (TGFbeta) superfamily members specify different functions: extracellular matrix interactions and growth factor bioavailability. J Biol Chem. 2011;286(7):5087–99.

9. Ling N, Ying SY, Ueno N, Esch F, Denoroy L, Guillemin R. Isolation and partial characterization of a Mr 32,000 protein with inhibin activity from porcine follicular fluid. Proc Natl Acad Sci U S A. 1985;82(21):7217–21.

10. Mason AJ, Farnworth PG, Sullivan J. Characterization and determination of the biological activities of noncleavable high molecular weight forms of inhibin A and activin A. Mol Endocrinol. 1996;10(9):1055–65.

11. Vale W, Rivier J, Vaughan J, McClintock R, Corrigan A, Woo W, et al. Purification and characterization of an FSH releasing protein from porcine ovarian follicular fluid. Nature. 1986;321(6072):776–9.

12. Ling N, Ying SY, Ueno N, Shimasaki S, Esch F, Hotta M, et al. A homodimer of the beta-subunits of inhibin A stimulates the secretion of pituitary follicle stimulating hormone. Biochem Biophys Res Commun. 1986;138(3):1129–37.

13. Mason AJ, Berkemeier LM, Schmelzer CH, Schwall RH. Activin B: precursor sequences, genomic structure and in vitro activities. Mol Endocrinol. 1989;3(9):1352–8.

14. Schang G, Ongaro L, Schultz H, Wang Y, Zhou X, Brule E, et al. Murine FSH Production Depends on the Activin Type II Receptors ACVR2A and ACVR2B. Endocrinology. 2020;161(7).

15. Rejon CA, Ho CC, Wang Y, Zhou X, Bernard DJ, Hebert TE. Cycloheximide inhibits follicle-stimulating hormone beta subunit transcription by blocking de novo synthesis of the labile activin type II receptor in gonadotrope cells. Cell Signal. 2013;25(6):1403–12.

16. Tzavlaki K, Moustakas A. TGF-beta Signaling. Biomolecules. 2020;10(3).

17. Fortin J, Boehm U, Deng CX, Treier M, Bernard DJ. Follicle-stimulating hormone synthesis and fertility depend on SMAD4 and FOXL2. FASEB J. 2014;28(8):3396–410.

18. Corpuz PS, Lindaman LL, Mellon PL, Coss D. FoxL2 Is required for activin induction of the mouse and human follicle-stimulating hormone beta-subunit genes. Mol Endocrinol. 2010;24(5):1037–51.

19. Lamba P, Fortin J, Tran S, Wang Y, Bernard DJ. A novel role for the forkhead transcription factor FOXL2 in activin A-regulated follicle-stimulating hormone beta subunit transcription. Mol Endocrinol. 2009;23(7):1001–13.

20. Tran S, Lamba P, Wang Y, Bernard DJ. SMADs and FOXL2 synergistically regulate murine FSHbeta transcription via a conserved proximal promoter element. Mol Endocrinol. 2011;25(7):1170–83.

21. Xu J, McKeehan K, Matsuzaki K, McKeehan WL. Inhibin antagonizes inhibition of liver cell growth by activin by a dominant-negative mechanism. J Biol Chem. 1995;270(11):6308–13.

22. Bernard DJ, Smith CL, Brule E. A Tale of Two Proteins: Betaglycan, IGSF1, and the Continuing Search for the Inhibin B Receptor. Trends Endocrinol Metab. 2020;31(1):37–45.

23. Brule E, Wang Y, Li Y, Lin YF, Zhou X, Ongaro L, et al. TGFBR3L is an inhibin B co-receptor that regulates female fertility. Sci Adv. 2021;7(51):eabl4391.

24. Li Y, Fortin J, Ongaro L, Zhou X, Boehm U, Schneyer A, et al. Betaglycan (TGFBR3) Functions as an Inhibin A, but Not Inhibin B, Coreceptor in Pituitary Gonadotrope Cells in Mice. Endocrinology. 2018;159(12):4077–91.

25. Lewis KA, Gray PC, Blount AL, MacConell LA, Wiater E, Bilezikjian LM, et al. Betaglycan binds inhibin and can mediate functional antagonism of activin signalling. Nature. 2000;404(6776):411–4.

26. Chapman SC, Bernard DJ, Jelen J, Woodruff TK. Properties of inhibin binding to betaglycan, InhBP/p120 and the activin type II receptors. Mol Cell Endocrinol. 2002;196(1-2):79–93.

27. Henen MA, Mahlawat P, Zwieb C, Kodali RB, Hinck CS, Hanna RD, et al. TGF-beta2 uses the concave surface of its extended finger region to bind betaglycan’s ZP domain via three residues specific to TGF-beta and inhibin-alpha. J Biol Chem. 2019;294(9):3065–80.

28. Wieteska L, Taylor AB, Punch E, Coleman JA, Conway IO, Lin YF, et al. Structures of TGF-beta with betaglycan and signaling receptors reveal mechanisms of complex assembly and signaling. Nat Commun. 2025;16(1):1778.

29. Goney MP, Wilce MCJ, Wilce JA, Stocker WA, Goodchild GM, Chan KL, et al. Engineering the Ovarian Hormones Inhibin A and Inhibin B to Enhance Synthesis and Activity. Endocrinology. 2020;161(8).

30. Walton KL, Kelly EK, Johnson KE, Robertson DM, Stanton PG, Harrison CA. A Novel, More Efficient Approach to Generate Bioactive Inhibins. Endocrinology. 2016;157(7):2799–809.

31. Pangas SA, Woodruff TK. Production and purification of recombinant human inhibin and activin. J Endocrinol. 2002;172(1):199–210.

32. Kappes EC, Kattamuri C, Czepnik M, Yarawsky AE, Brule E, Wang Y, et al. Follistatin Forms a Stable Complex With Inhibin A That Does Not Interfere With Activin A Antagonism. Endocrinology. 2023;164(3).

33. Goehring A, Lee CH, Wang KH, Michel JC, Claxton DP, Baconguis I, et al. Screening and large-scale expression of membrane proteins in mammalian cells for structural studies. Nat Protoc. 2014;9(11):2574–85.

34. Lichty JJ, Malecki JL, Agnew HD, Michelson-Horowitz DJ, Tan S. Comparison of affinity tags for protein purification. Protein Expr Purif. 2005;41(1):98–105.

35. Yeliseev A, Zoubak L, Schmidt TGM. Application of Strep-Tactin XT for affinity purification of Twin-Strep-tagged CB(2), a G protein-coupled cannabinoid receptor. Protein Expr Purif. 2017;131:109–18.

36. Witte CP, Noel LD, Gielbert J, Parker JE, Romeis T. Rapid one-step protein purification from plant material using the eight-amino acid StrepII epitope. Plant Mol Biol. 2004;55(1):135–47.

37. Bolanos-Garcia VM, Davies OR. Structural analysis and classification of native proteins from E. coli commonly co-purified by immobilised metal affinity chromatography. Biochim Biophys Acta. 2006;1760(9):1304–13.

38. Makanji Y, Temple-Smith PD, Walton KL, Harrison CA, Robertson DM. Inhibin B is a more potent suppressor of rat follicle-stimulating hormone release than inhibin a in vitro and in vivo. Endocrinology. 2009;150(10):4784–93.

39. Schmidt TG, Koepke J, Frank R, Skerra A. Molecular interaction between the Strep-tag affinity peptide and its cognate target, streptavidin. J Mol Biol. 1996;255(5):753–66.

40. Schmidt TG, Skerra A. The Strep-tag system for one-step purification and high-affinity detection or capturing of proteins. Nat Protoc. 2007;2(6):1528–35.

41. Voss S, Skerra A. Mutagenesis of a flexible loop in streptavidin leads to higher affinity for the Strep-tag II peptide and improved performance in recombinant protein purification. Protein Eng. 1997;10(8):975–82.

42. Abramson J, Adler J, Dunger J, Evans R, Green T, Pritzel A, et al. Accurate structure prediction of biomolecular interactions with AlphaFold 3. Nature. 2024;630(8016):493–500.

43. Kapust RB, Tozser J, Copeland TD, Waugh DS. The P1’ specificity of tobacco etch virus protease. Biochem Biophys Res Commun. 2002;294(5):949–55.

44. Fan X, Li X, Zhou Y, Mei M, Liu P, Zhao J, et al. Quantitative Analysis of the Substrate Specificity of Human Rhinovirus 3C Protease and Exploration of Its Substrate Recognition Mechanisms. ACS Chem Biol. 2020;15(1):63–73.

45. Tillet E, Ouarne M, Desroches-Castan A, Mallet C, Subileau M, Didier R, et al. A heterodimer formed by bone morphogenetic protein 9 (BMP9) and BMP10 provides most BMP biological activity in plasma. J Biol Chem. 2018;293(28):10963–74.

46. Cheifetz S, Bassols A, Stanley K, Ohta M, Greenberger J, Massague J. Heterodimeric transforming growth factor beta. Biological properties and interaction with three types of cell surface receptors. J Biol Chem. 1988;263(22):10783–9.

47. Israel DI, Nove J, Kerns KM, Kaufman RJ, Rosen V, Cox KA, et al. Heterodimeric bone morphogenetic proteins show enhanced activity in vitro and in vivo. Growth Factors. 1996;13(3-4):291–300.

48. Ling N, Ying SY, Ueno N, Shimasaki S, Esch F, Hotta M, et al. Pituitary FSH is released by a heterodimer of the beta-subunits from the two forms of inhibin. Nature. 1986;321(6072):779–82.

49. Montague TG, Schier AF. Vg1-Nodal heterodimers are the endogenous inducers of mesendoderm. Elife. 2017;6.

50. Derynck R, Budi EH. Specificity, versatility, and control of TGF-beta family signaling. Sci Signal. 2019;12(570).

51. Heldin CH, Moustakas A. Signaling Receptors for TGF-beta Family Members. Cold Spring Harb Perspect Biol. 2016;8(8).

52. Miller DSJ, Schmierer B, Hill CS. TGF-beta family ligands exhibit distinct signalling dynamics that are driven by receptor localisation. J Cell Sci. 2019;132(14).

53. Little SC, Mullins MC. Bone morphogenetic protein heterodimers assemble heteromeric type I receptor complexes to pattern the dorsoventral axis. Nat Cell Biol. 2009;11(5):637–43.

54. Kim SK, Whitley MJ, Krzysiak TC, Hinck CS, Taylor AB, Zwieb C, et al. Structural Adaptation in Its Orphan Domain Engenders Betaglycan with an Alternate Mode of Growth Factor Binding Relative to Endoglin. Structure. 2019;27(9):1427–42 e4.

55. Schindelin J, Arganda-Carreras I, Frise E, Kaynig V, Longair M, Pietzsch T, et al. Fiji: an open-source platform for biological-image analysis. Nat Methods. 2012;9(7):676–82.

