## Supplemental figures for "An efficient system for production of highly purified inhibins using mammalian cell expression and tandem affinity purification"

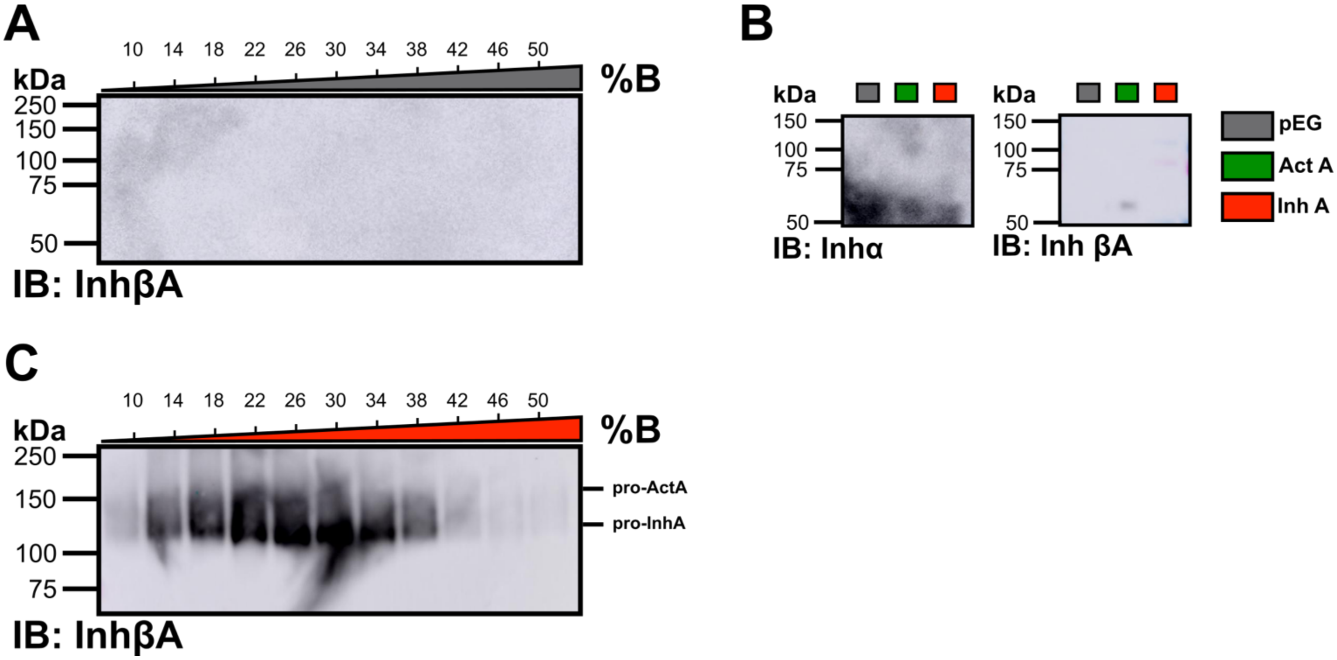


**Figure S1: Expi293 cells do not produce ActA or InhA endogenously. (A)** Conditioned medium (CM) corresponding to expi293 cells transfected with the empty vector (pEG) was loaded onto the Co-IMAC column, and eluted fractions (0.55 μL/lane) were immunoblotted against the InhβA subunit. **(B)** Immunoblots of combined flow-through and wash (0.55 μL/lane) from the Co-IMAC purifications of CM from expi293 cells transfected with the empty vector (pEG), InhβA V1 construct (ActA V1), or InhA V1 construct. **(C)** Eluted Co-IMAC fractions (0.55 μL/lane) from the purification of WT InhA (V3 construct) were run longer to obtain better separation between pro-ActA and pro-InhA prior to immunoblotting with the inhβA subunit antibody.

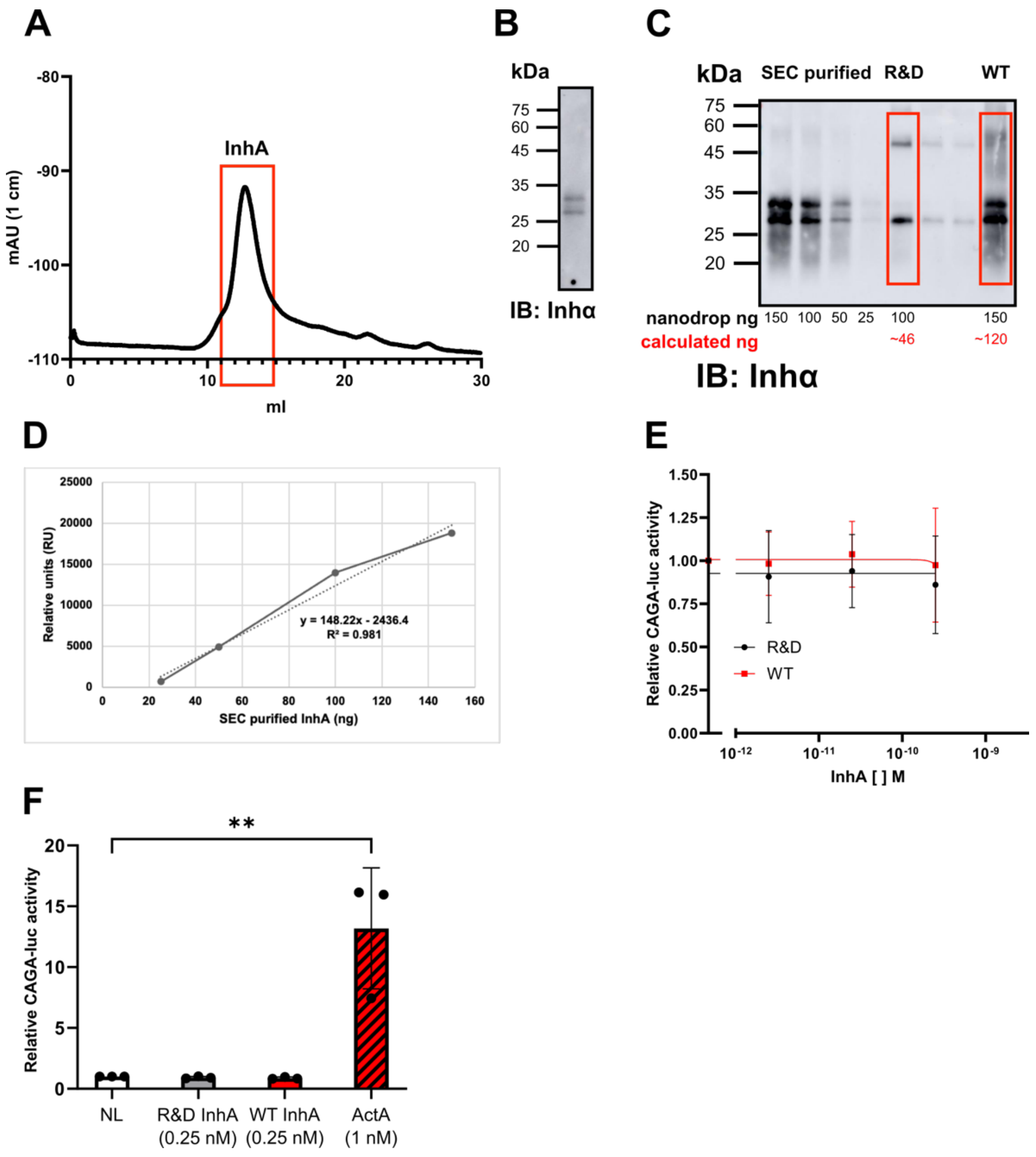


**Figure S2: Size exclusion chromatography (SEC) purified InhA can be used to estimate the concentration of mature GF in inhibin preparations. (A)** InhA was purified by SEC, and **(B)** assessed by immunoblotting using the inhα antibody. **(C)** The indicated amount of SEC purified InhA, InhA from R&D, and our WT InhA were immunoblotted against the inhα subunit. The amount of mature GF calculated from the standard curve in **(D)** is indicated in red. CHO cells were transfected with the CAGA-luc promoter-reporter and pCMV5 (empty vector). In **(E)**, cells were treated with 1 nM ActA (R&D) and the indicated concentration of InhA from R&D or our non-SEC purified InhA. Reporter activity was normalized to ActA alone. Note, betaglycan ZPC is not included in this analysis. In **(E)**, data represent mean values (error bars depicting SD), and were analyzed by two-way ANOVA, followed by Sidak's multiple comparisons test (n = 4 experimental replicates). **(F)** Cells were treated with the indicated amount of ligand (the maximal amount of InhA or ActA used in **Fig. 7C**), and responses were normalized to the no-ligand (NL) condition. In **(F)**, data are presented as individual values with the height of the bar graph depicting the mean (error bars depicting SD) and analyzed by one-way unpaired t-test (n = 3 experimental replicates, each point is an individual replicate; ** P < 0.01)

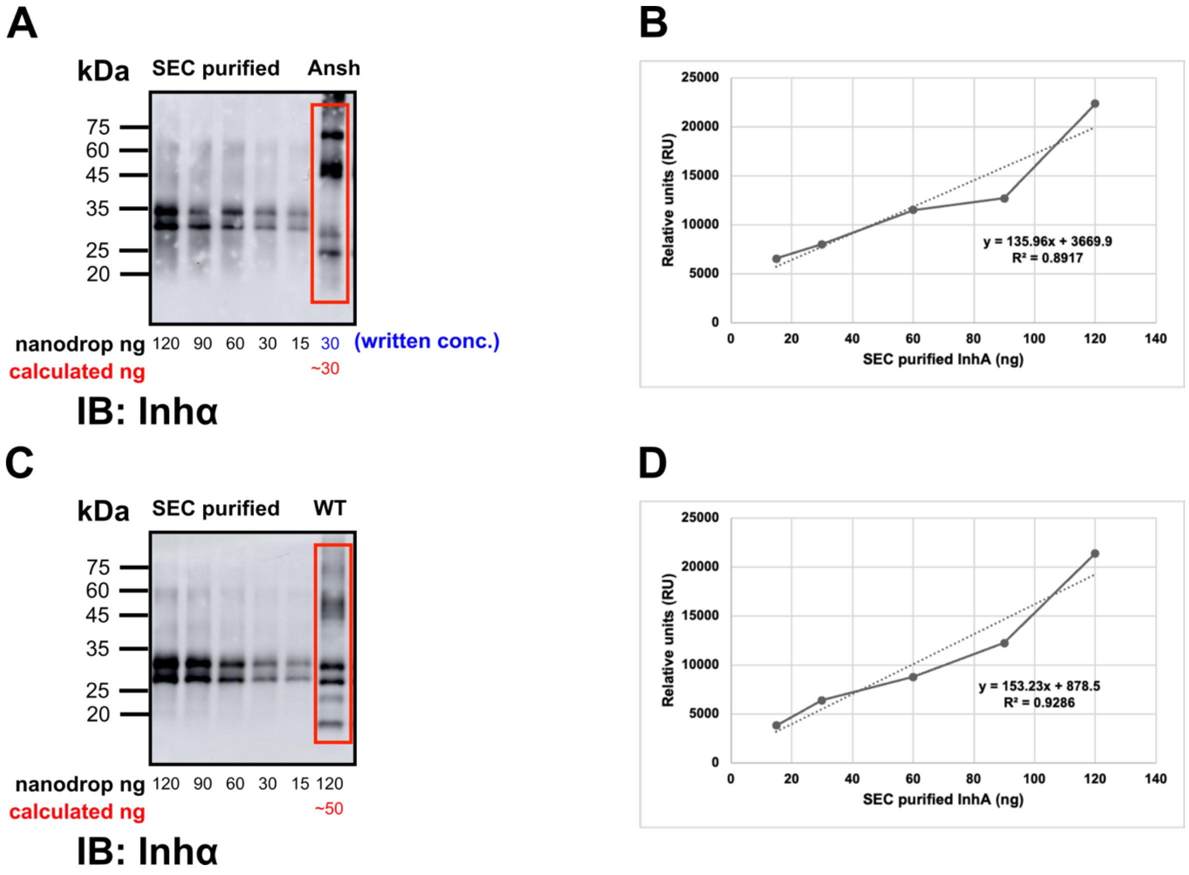


**Figure S3: SEC purified InhA can be used to infer the amount of mature GF in InhB preparations. (A)** To determine the amount of mature GF in Anshlabs InhB, SEC purified InhA and Anshlabs InhB were immunoblotted against the inhα subunit. The amount of mature GF as determined from the standard curve of SEC purified InhA **(B)** is written below in red. The same analysis was conducted with our InhB **(C-D)**.

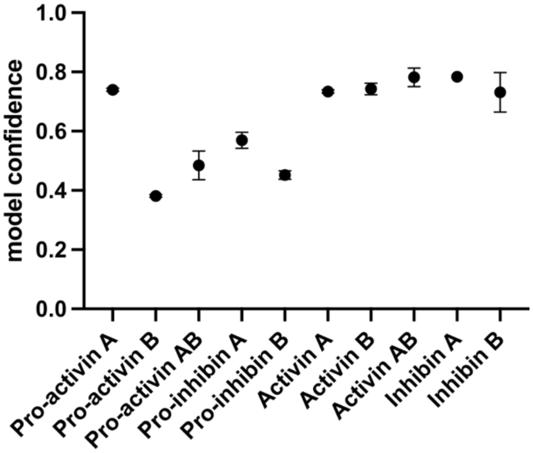


**Figure S4: The confidence of AlphaFold3 predicted structures of activin and inhibin decreases with the inclusion of pro-inhβB.** AlphaFold3 model confidence scores (0.8*iPTM + 0.2*PTM) for activins and inhibins, with and without their associated pro-domains (error bars depicting SD, n = 3, individual predictions).
